# Understanding the emergence of contingent and deterministic exclusion in multispecies communities

**DOI:** 10.1101/2020.09.23.310524

**Authors:** Chuliang Song, Lawrence H. Uricchio, Erin A. Mordecai, Serguei Saavedra

## Abstract

Competitive exclusion can be classified as deterministic or as historically contingent. While competitive exclusion is common in nature, it has remained unclear when multispecies communities should be dominated by deterministic or contingent exclusion. Here, we provide a general theoretical approach to explain both the emergence and sources of competitive exclusion in multispecies communities. We illustrate our approach on an empirical competition system between annual and perennial plant species. First, we find that the life-history of perennial species increases the probability of observing contingent exclusion by increasing their effective intrinsic growth rates. Second, we find that the probability of observing contingent exclusion increases with weaker intraspecific competition, and not with the level of hierarchical competition. Third, we find a shift from contingent exclusion to dominance with increasing numbers of competing species. Our work provides a heuristic framework to increase our understanding about the predictability of species persistence within multispecies communities.

## Introduction

Species coexistence is one of the most studied topics in ecology (Vellend, 2016); however, some have observed that competitive exclusion is the norm rather than the exception in nature (Hardin, 1960; Goldford *et al*., 2018; Blowes *et al*., 2019). Indeed, coexisting species within ecological communities are usually a fraction of all the species available in a local species pool (Odum *et al*., 1971; Sigmund, 1995). Exclusion as a ubiquitous feature of ecological communities has been demonstrated empirically across a wide range of life forms, including algae (Narwani *et al*., 2013), annual plants (Godoy & Levine, 2014a), microbiomes (Friedman *et al*., 2017), bacteria (Tan *et al*., 2017), and nectar-colonizing yeasts (Grainger *et al*., 2019). Importantly, due to the inherent stochasticity in community assembly, competitive exclusion can be broadly classified into two ecologically different categories (Fukami, 2015; Grainger *et al*., 2019). One category is *deterministic exclusion* (also known as dominance). That is, the order of species arrivals does not affect which species is competitively excluded. The other category is *contingent exclusion* (also known as priority effects). That is, the order of species arrivals does affect which species is competitively excluded. Knowing whether competitive exclusion is deterministic or contingent is fundamental to understanding the role of predictability and randomness in community assembly (Lawton, 1999; Fukami, 2015). For example, it has direct implications for conservation management: depending on whether the exclusion of native species is deterministic or contingent, we should adopt different strategies to restore biodiversity resulting after exotic species invasion (Bøhn *et al*., 2008; McGeoch *et al*., 2016).

Since the 1930s, theoretical and empirical research has systematically documented and expanded our understanding of competitive exclusion between two competing species (Gause, 1932; Ayala, 1969; Brown, 1971; Gilpin & Justice, 1972). Moreover, in recent decades, theoretical studies have started to provide an overarching framework to synthesize data across different competition systems (Mordecai, 2013; Johnson & Bronstein, 2019; Ke & Wan, 2020). This theoretical development started by focusing on the conditions leading to deterministic exclusion (Chesson, 2000; Adler *et al*., 2007), and then it was extended to investigate the conditions for contingent exclusion (Mordecai, 2011; Fukami *et al*., 2016; Ke & Letten, 2018). Similarly, extensive empirical research started to examine the sources of deterministic exclusion (Mayfield & Levine, 2010; Violle *et al*., 2011; Adler *et al*., 2010), and more recently it has moved to the analysis of contingent exclusion (Grainger *et al*., 2018, 2019; Song *et al*., 2020a). Focusing on competition between two species, this body of work has shown that deterministic exclusion is more likely to occur when the competitively inferior species has a lower intrinsic growth rate and when intraspecific interactions are stronger than interspecific interactions. By contrast, greater similarity in species intrinsic growth rates and stronger interspecific relative to intraspecific interactions promote contingent exclusion (Ke & Letten, 2018; Song *et al*., 2020a).

However, it remains unclear whether these clear conditions at the *two-species* level also operate in *multispecies* communities. First, the aforementioned body of work has been mainly executed under a theoretical formalism for two-species communities, which does not have a counterpart for multispecies communities. Specifically, the standard formalism for two-species communities is incompatible with the current canonical formalism for multispecies communities (Song *et al*., 2019). While the formalism for two-species communities can easily distinguish competitive exclusion into deterministic exclusion and contingent exclusion, the formalism for multispecies communities cannot distinguish them as easily (Barabás *et al*., 2018). Second, the patterns of contingent and deterministic exclusion are inherently more complicated in multispecies communities. For example, multispecies communities may exhibit a mixed outcome of competitive exclusion: some species can be deterministically excluded while others can be contingently excluded. This implies that we cannot always classify the competition dynamics of a community simply as either deterministic or contingent in multispecies communities, which is typically done in two-species communities. Instead, competitive exclusion in multispecies communities should be analyzed at the species level. Specifically, for a community with *S* interacting species, there are in total *S*! possibilities of species arrival orders. We classify competitive exclusion as follows: if a species is competitively excluded in all possible arrival orders, then the species is deterministically excluded; if a species is competitively excluded in some but not all possible arrival orders, then the species is contingently excluded. Thus, we still lack a full understanding about competitive exclusion in species-rich ecological communities, where more complex dynamics, including non-hierarchical competition and higher-order interactions, can occur (Levine *et al*., 2017; Saavedra *et al*., 2017).

The complexity of competitive exclusion in multispecies communities calls for further developing the existing theory or establishing new approaches. In this line, the *structural approach* in ecology has provided an alternative theoretical perspective to study competitive exclusion in multispecies communities (Saavedra *et al*., 2017; Song *et al*., 2018b). In general, the structural approach posits that how likely a particular outcome of competition is to occur can be understood through the full range of environmental conditions (contexts) compatible with that qualitative outcome. While the structural approach was initially devised to investigate species coexistence as the qualitative outcome (Rohr *et al*., 2014; Saavedra *et al*., 2017), it can also be extended to study competitive exclusion (Song *et al*., 2020a). Here, we apply the structural approach to investigate the emergence and sources of competitive exclusion in multispecies communities as a function of species’ intrinsic growth rates, community size (number of competing species), and competition structure (i.e., the interaction matrix).

As an empirical application for our framework, we use data on five grass species from California grasslands. The invasion of exotic annual species presumably has, together with human-induced habitat shifts, competitively excluded native perennial species in many regions. This has been considered as “one of the most dramatic ecological invasions worldwide” (Seabloom *et al*., 2003). Indeed, empirical evidence suggests that long-term, stable coexistence of multiple annual and perennial species is unlikely (Uricchio *et al*., 2019). However, most theoretical (Crawley & May, 1987; Rees & Long, 1992; Kisdi & Geritz, 2003; Uricchio *et al*., 2019) and experimental studies (Hamilton *et al*., 1999; Corbin & D’Antonio, 2004; Seabloom *et al*., 2003; Mordecai *et al*., 2015) have primarily focused on the competitive exclusion between two species (i.e., one annual species and one perennial species). Thus, it remains unclear how these ecological dynamics are expected to play out among multiple annual and perennial species. To this end, we apply our investigation to field experiments on three exotic annual species (*Bromus hordeaceus, Bromus diandrus*, and *Avena barbata*) and two native perennial species (*Elymus glaucus* and *Stipa pulchra*) that occur in California grasslands (Uricchio *et al*., 2019). Previous simulation-based work showed a complex pattern of coexistence, deterministic exclusion, and contingent exclusion among these species (Uricchio *et al*., 2019). In addition, competition among these species is intransitive (non-hierarchical), and stronger between species than within species (i.e., self-regulation is weak). Here, we integrate a structural approach with numerical simulations to systemically disentangle the contributions of life-history traits, community size, and competition structure to deterministic and contingent exclusion in California grasslands.

## Methods

### Structural approach to competitive exclusion

The structural approach in ecology is built on a systematic and probabilistic understanding of how likely a given type of qualitative dynamics is to occur (Song, 2020; Saavedra *et al*., 2020). Here, the qualitative dynamics of interest are deterministic exclusion and contingent exclusion. The structural approach simplifies ecological dynamics as a function of internal and external conditions (Saavedra *et al*., 2017). External conditions are phenomenologically represented by *intrinsic growth rates* (the maximum growth rate a species can have in isolation) and they are assumed to change in response to environmental conditions. Internal conditions are phenomenologically represented by the *competition structure* (the matrix whose elements correspond to the competitive effect of one species on another) and are assumed to be fixed across time (see Appendix B for an in-depth discussion). This characterization and set of assumptions allows us to calculate the domain of external conditions (the context) compatible with a given qualitative outcome as a function of a given set of internal conditions. The larger this domain is, the higher the probability that the observed external conditions match with one inside the domain, leading to the realization of the corresponding qualitative outcome.

Formally, the structural approach uses the *feasibility domain* as the domain of external conditions compatible with a given qualitative outcome. The feasibility domain describes the full range of intrinsic growth rates compatible with positive abundances of all species in the community (i.e., feasible equilibrium). While the competition structure determines the *shape* of the feasibility domain (Song *et al*., 2018b, 2020a; Tabi *et al*., 2020), the observed intrinsic growth rates determine whether the community is inside or outside of the feasibility domain (Saavedra *et al*., 2017). When the community is outside of the feasibility domain, the community is expected to be driven by deterministic exclusion. To further understand the qualitative dynamics when the community is inside the feasibility domain, we need to consider the *orientation* of the feasibility domain in addition to its shape. The orientation refers to whether the feasible equilibrium in the feasibility domain is dynamically stable or not. The importance of the orientation is that stable feasibility leads to coexistence, whereas unstable feasibility leads to contingent exclusion (Case, 1999; Fukami *et al*., 2016). The orientation of the feasibility domain is mainly driven by the ratio of intra-to interspecific interactions (Song *et al*., 2020a). In sum, following the structural approach, whether competitive exclusion is deterministic or contingent should be expected to be mainly driven by the match between the observed intrinsic growth rates (mainly constrained by life-history processes) with the shape and the orientation of the feasibility domain (both of which are determined by the observed competition structure). Note that our framework is only an expectation given that multispecies dynamics is a function of the underlying complexity of a system (AlAdwani & Saavedra, 2020).

By way of example, focusing on two-species communities (see Figure 1 for a graphical illustration), one can establish three key intuitions about competitive exclusion derived from the structural approach (Song *et al*., 2020a): (i) For contingent exclusion to occur, it is necessary that species depress their competitor’s per capita growth rate more than their own (changing the orientation of the feasibility domain). (ii) The larger the intrinsic growth rate of the competitively inferior species, the more likely contingent exclusion is to occur. (iii) The larger the feasibility domain, the more likely contingent exclusion is to occur. Note that these intuitions are aligned with the theoretical expectations from frameworks based on growth rates when rare that are explicitly justified for two-species communities (Adler *et al*., 2007; Fukami *et al*., 2016). We hypothesize these three intuitions operate in multispecies communities as heuristic rules, which we test in the empirical dataset. It is worth noting that on average, the size of the feasibility domain decreases with the number of species in a community (Grilli *et al*., 2017; Song *et al*., 2018b). Thus, following these premises, contingent exclusion should be more likely to occur in ecological communities (i) with fewer number of species, (ii) with species that more strongly depress their competitor’s growth rate relative to their self-regulation, and (iii) where life-history processes increase the intrinsic growth rates of competitively inferior species.

**Figure 1:**
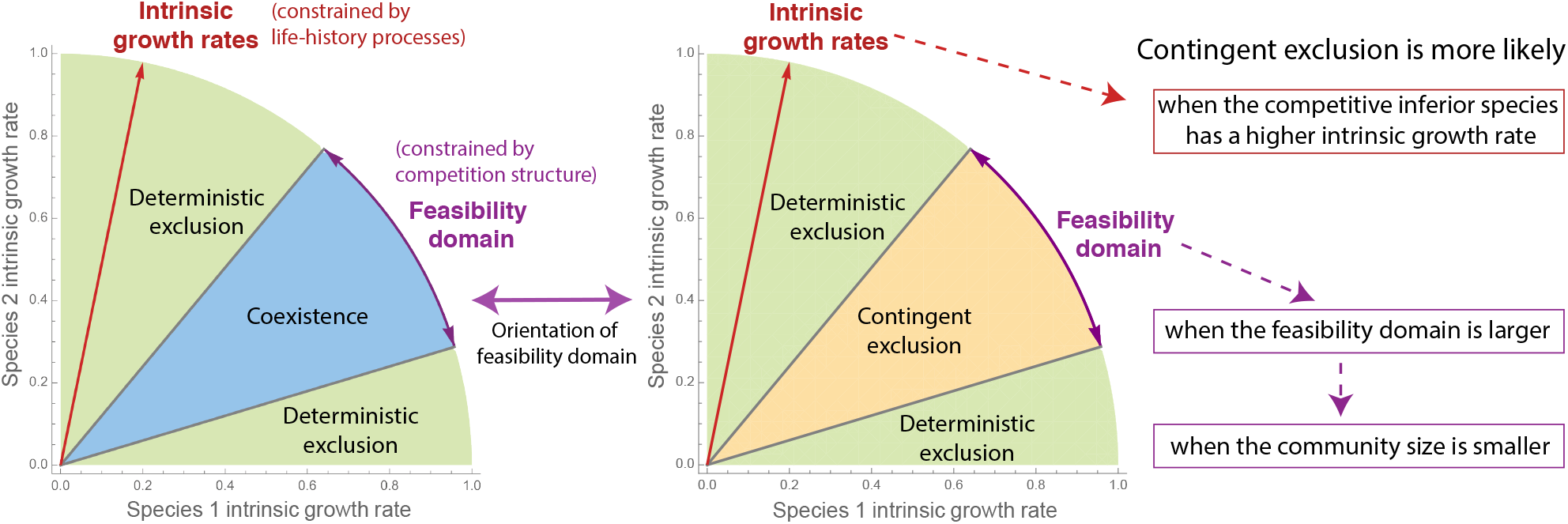
Three key intuitions on competitive exclusion following a structural approach. For a hypothetical community with two competing species, the figure shows the parameter space defined by the intrinsic growth rates (phenomenological abiotic conditions) of the two species. The feasibility domain (middle blue or orange region) is the set of all directions of intrinsic growth rates compatible with a feasible equilibrium. If the feasible equilibrium is dynamically unstable, the region corresponds parameters that are compatible with contingent exclusion (right panel: orange region); if the feasible equilibrium is dynamically stable, the region is compatible with stable coexistence (left panel: blue region). The complement of the feasibility domain regardless of dynamical stability (green region) corresponds to the directions of intrinsic growth rates associated with deterministic exclusion. Following the structural approach in ecology, we can derive three key intuitions: (i) Contingent exclusion is expected to be more likely when the competitive inferior species has a higher intrinsic growth rate. (ii) Contingent exclusion is more likely when the feasibility domain is larger. (iii) Contingent exclusion is more likely when the community size is smaller. The ecological rationale is that adding a new species generally further constrains the feasibility domain to be smaller. Note that the third intuition is a corollary from the second intuition since the feasibility domain generally shrinks with community size.

### Population dynamics of annual and perennial species

To study ecological dynamics under a structural approach, it is necessary to assume the governing laws of population dynamics (Cenci & Saavedra, 2018). Annual and perennial species have different population dynamics. A key difference is that annual species only carry over between growing seasons as seeds, while perennial species carry over between growing seasons as both seeds and adults. To simplify the notation, for each species *i* we hereafter denote annual seeds as *N*_*i*_, perennial seeds as 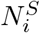, and perennial adults as 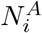.

Focusing on annual species, we assume the classic seed-banking annual plant model with Beverton-Holt competition (Levine & HilleRisLambers, 2009; Godoy & Levine, 2014b). For annual plants, these dynamics can be written as (illustrated in Figure 2A)

**Figure 2:**
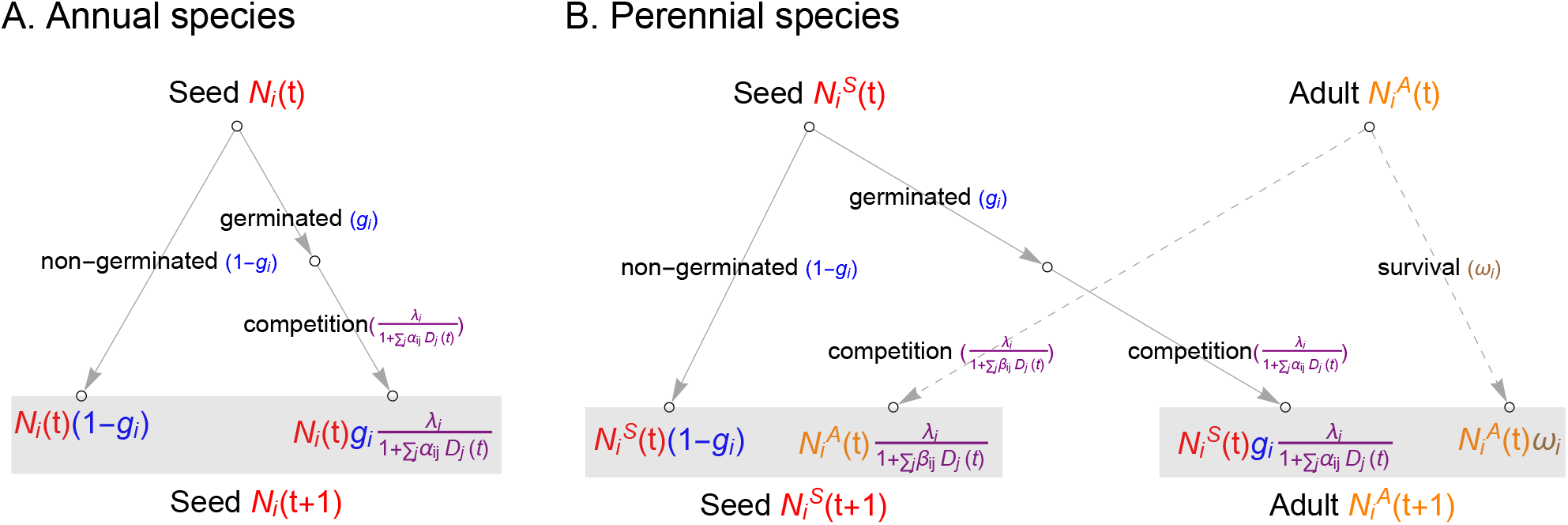
Population dynamics of annual and perennial plant species. Panel (A) illustrates the population dynamics of an annual plant species (Eqn. 1). Annual plant dynamics are tracked as seeds entering each growing season. Some annual seeds germinate, and the germinated seeds produce seeds at a rate reduced by competition from other plant species. Panel (B) illustrates the dynamics of a perennial plant species (Eqn. 3 and 4). The perennial plant has two life stages, seed and adult. Some perennial seeds germinate, and the germinated seeds would produce adults at a rate reduced by competition from other plant species (left side). Perennial life history: some perennial adults survive as perennials, while some perennial adults produce seeds and are decreased by competition from other plant species (right side, dashed lines). Note that the dynamics of perennial plants can be be modeled with or without these perennial life-history processes.

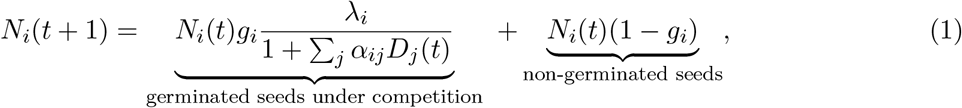

where *N*_*i*_ is the number of seeds of species *i, g*_*i*_ is the germination fraction, *λ*_*i*_ is per-capita seed production in the absence of competition, and *α*_*ij*_ is the per-capita competitive effect of species *j* on species *i*. The summation of the germinated density *D*_*j*_ is established over all species of annual germinants, perennial germinants, and perennial adults. Specifically, the germinated density *D*_*j*_ of competitors from species *j* is

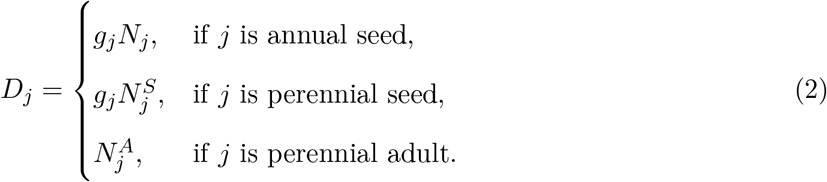

Perennial seed population dynamics can be written as (illustrated in Figure 2B)

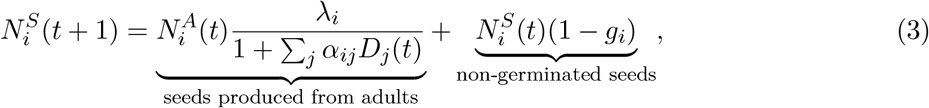

which is a slight modification of the annual plant model. Specifically, perennial seeds are generated when adults *A*_*i*_ reproduce, and reduced by both species competition (first term in Eqn. 3) and the survival of non-germinating perennial seeds (second term in Eqn. 3). The competition coefficients *α*_*ij*_ and densities *D*_*j*_ are defined as above (Eqn. 2).

Finally, the population dynamics of perennial adults can be written as (illustrated in Figure 2B)

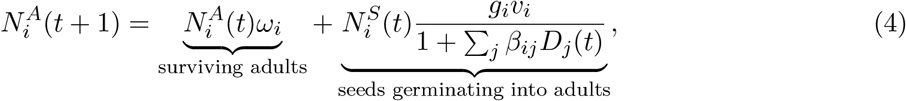

where *ω*_*i*_ is the over-summer survival fraction of perennial adults, and *v*_*i*_ is the fraction of over-summer maturation from perennial seedlings into adults for the following year (in the absence of competition). Note that perennial adults are generated by both surviving perennial adults *A*_*i*_ (first term in Eqn. 4) and seeds *S*_*i*_ that germinate and survive over the summer to become adults. Again, the abundance of perennial adults are reduced by species competition (second term in Eqn. 4).

### Empirical data and patterns of competitive exclusion

We based our analysis on an experimental study conducted in 2015-2016 in Jasper Ridge Biological Preserve, located in San Mateo County, California (377°24’N, 122°13’30”W; 66–207 m) (Uricchio *et al*., 2019). The experimental study investigated five focal grassland species with three exotic annual species (*Avena barbata, Bromus diandrus*, and *Bromus hordeaceus*) and two native perennial species (*Stipa pulchra* and *Elymus glaucus*). These species were studied because they were abundant and widespread in California grasslands. This experimental study measured key demographic rates that determined species growth, including seed overwinter survival, germination, establishment, adult bunchgrass survival, and the effects of competition on per-capita seed production (Uricchio *et al*., 2019). In addition, the study measured competition experimentally and observationally in 1-*m*^2^ plots. This covered a broad range of naturally occurring plant densities. Competition and growth parameters were sampled via Markov-Chain Monte Carlo based on population dynamics models developed for the three annual and two perennial grass species. We used 2000 samples from the joint posterior distribution of these parameters to conduct our study.

Given the timescale of competitive exclusion in natural grassland communities, the empirical study did not perform experiments on competitive exclusion. Thus, we employ the experimentally-parameterized population dynamics of annual and perennial species to simulate the patterns of competitive exclusion. Specifically, for a community with *S* interacting species, we simulate all *S*! possible species arrival orders. Each species arrives into the community when the community has already reached its stationary state, and we focus on the final stationary state. Using the final stationary states across all arrival orders we can classify a species as either contingently excluded (excluded in some arrival orders), deterministically excluded (excluded in all arrival orders), or persistent (not excluded in any arrival orders). Importantly, note that the classification of species is based solely on the dynamical outcomes derived from numerical simulations, which is not directly related to whether the community is feasible or dynamically stable (AlAdwani & Saavedra, 2020). This also prevents a tautological link between the classification scheme and the structural approach.

### Understanding the sources of competitive exclusion

To understand the emergence of deterministic and contingent exclusion, it is necessary to understand their sources. For this purpose, here we focus on three key ecological properties: life-history processes, community size, and competition structure. Following a structural approach, we investigate these three sources in the California grassland study system.

#### Life-history processes

Annual and perennial species differ in their strategies for persisting between growing seasons, either solely as seeds or additionally as surviving adults (Lundgren & Des Marais, 2020)—as we have exemplified in our population dynamics models. To understand the contribution of this life-history difference to the emergence of competitive exclusion, we applied the structural approach to the population dynamics of species with and without modeling the life-history difference between annual and perennial species.

By removing over-summer survival of adult perennials and assuming that germinating seeds produce new seeds within the same growing season, thereby removing the life-history difference between annual and perennial species (i.e., removing the dashed links in Figure 2B), the feasibility condition of a multispecies community reduces to

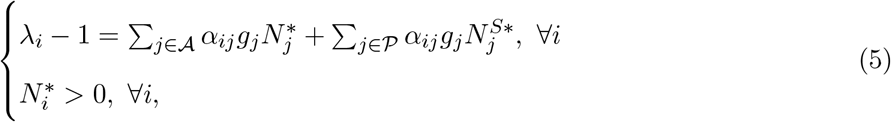

where 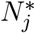 represents either the annual or the perennial species, *𝒜* represents the set of all annual species, and *𝒫* represents the set of all perennial species.

Alternatively, incorporating the life-history processes of perennial species (i.e., keeping the dashed links in Figure 2B), the feasibility condition is

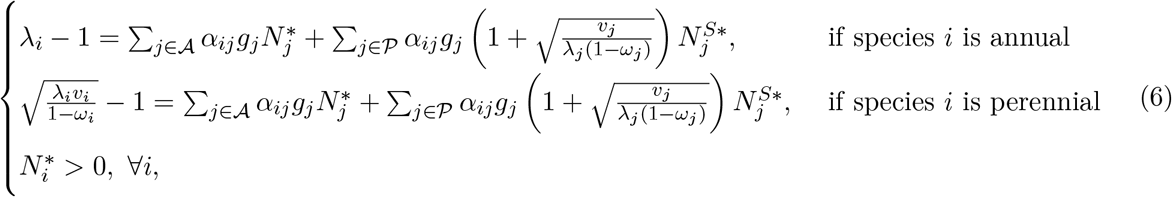

where again 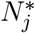 represents either the annual or the perennial species, *𝒜* represents the set of all annual species, and *𝒫* represents the set of all perennial species. The derivations can be found in Appendix C.

Importantly, the feasibility domain of the multispecies communities is the same excluding (Eqn. 5) or including (Eqn. 6) perennial life-history processes. The mathematical rationale of this identity comes from the column scaling invariance of the feasibility domain (Song *et al*., 2020b) (Appendix E). The ecological rationale can be interpreted by the fact that perennial life-history processes affect only the absolute equilibrium abundances, and not the competition coefficients (Saavedra *et al*., 2017). Thus, for the assumed population dynamics, the feasibility domain of the multispecies community is uniquely determined by the competition structure {*a*_*ij*_} summarized in the interaction matrix, but not by any other parameter. This result additionally implies that life-history processes only affect the patterns of competitive exclusion (whether it is dominated by deterministic or contingent exclusion) by changing the effective intrinsic growth rates. Specifically, life-history processes change the effective intrinsic growth rates of the perennial species from (*λ*_*i*_ − 1) to 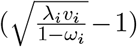 (see Appendix C for variations of assumptions).

We test the effects of life history differences on competitive exclusion in the species present in our empirically parameterized California grassland system. As we show theoretically, the effects can only come through the effective intrinsic growth rates. It is unclear *a priori* whether the life-history processes increase or decrease the effective intrinsic growth rates of the perennial species empirically.

#### Community size

As described above, following a structural approach, deterministic exclusion is expected to dominate over contingent exclusion in species-rich communities (see section *Structural approach on competitive exclusion*, Figure 1). In order to investigate the contribution of community size to the patterns of competitive exclusion, we need to analyze how the probabilities of observing deterministic and contingent exclusion for each species change as a function of community size. Importantly, while the theory suggests that we should get more deterministic exclusion as community size increases, it is possible that the observed parameters from empirical communities do not support this pattern. Here we test whether these theoretical patterns hold in the California grassland system.

#### Competition structure

Ecological communities are characterized by non-random competition structures (Thébault & Fontaine, 2010; Song *et al*., 2018a; Song & Saavedra, 2020). Indeed, Figure 5A shows the inferred competition structure (the direction and strength of species competition) of annual and perennial species in the California grassland system. This figure reveals two key features of the empirically studied competition structure. First, the intraspecific competition (self-regulation) is generally weaker than the interspecific competition. Second, interspecific competition forms an intransitive structure (also known as a non-hierarchical structure). The importance of these two features has been a central question in ecological research (Soliveres *et al*., 2015; Gallien *et al*., 2017; Barabás *et al*., 2017; Kinlock, 2019).

To test the overall effect of the competition structure on the patterns of competitive exclusion, we investigate how the competition structure changes the size of the feasibility domain in the empirical parameter space estimated for California grassland species. Recall that it is expected that contingent exclusion dominates multispecies communities with larger feasibility domains. We compute numerically the size of the feasibility domain from Eqn. (6) (Song *et al*., 2018b). Additionally, to separate the specific contributions of the two structural features of competition (i.e., intraspecific competition and intransitive competition), we use model-generated communities with four types of competition structures: (i) communities with either weak (intraspecific*<*interspecific) or strong (intraspecific>interspecific) intraspecific competition, and (ii) communities with either a hierarchical or intransitive competition structure. Focusing on the first structural combination, we consider strong intraspecific competition when the intraspecific competition of a given species is larger than the sum of the interspecific competition that this species experiences from other species (the opposite for weak intraspecific competition). Focusing on the second structural combination, we first generate a Erdős-Rényi structure as an instrumental initiation where each competition strength is independently sampled from a uniform distribution [0, 1](Song & Saavedra, 2018), and then we arrange the competition structure as either hierarchical or intransitive. We investigate which combinations can reproduce the associations between competitive exclusion and feasibility domain observed in the empirical data. We have tested other parameterizations to evaluate the robustness (Appendix F).

## Results

We first analyzed the effects of perennial life-history processes on whether a community is dominated by deterministic or contingent exclusion. The structural approach postulates that contingent exclusion is more likely when competitively inferior species have higher intrinsic growth rates (Figure 1). Theoretically, perennial life-history processes only regulate the intrinsic growth rates—via their effects on survival and fecundity in the absence of competition—but not the feasibility domain, which exclusively depends on competition structure. Because the perennial species included in this study were generally competitively inferior to the annual species, we expected that incorporating perennial life-history processes would yield a higher frequency of contingent exclusion by increasing perennial species intrinsic growth rates.

Focusing on all possible two-species communities with one annual and one perennial species, Figure 3 confirms the expectation that perennial life-history processes promote contingent exclusion. To illustrate this effect, we used a standard graphical representation of ecological dynamics for two species: the niche-overlap-fitness-ratio space (Adler *et al*., 2007; Chesson & Kuang, 2008). Specifically, Figure 3 shows that by adding perennial life-history processes to the model, the species average fitness of perennial species increases, which leads to a higher frequency of contingent exclusion, rather than deterministic exclusion. In addition, we found that incorporating life-history processes can change the outcome of the dynamics when subject to different types of environmental perturbations acting on parameters (Song *et al*., 2020a). That is, we found that communities exhibit robustness to perturbations acting on intrinsic growth rates but not on competition strength when perennial life-history is excluded, while they exhibit robustness to perturbations acting on competition strength but not on intrinsic growth rates when perennial life-history is incorporated (see Appendix D). Importantly, multispecies communities exhibit qualitatively identical patterns (see Figure 4).

**Figure 3:**
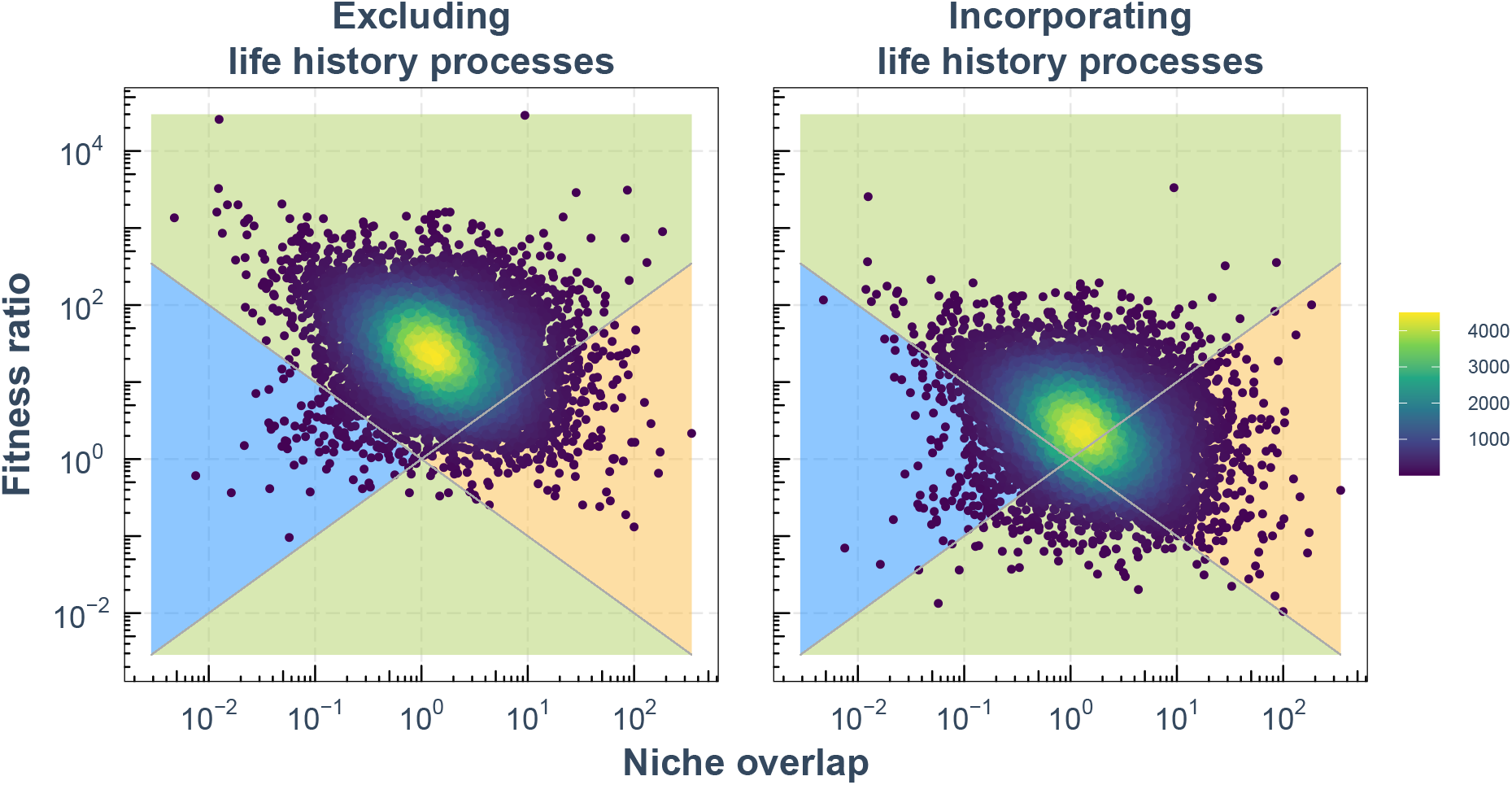
Perennial life-history processes increase the frequency of contingent exclusion by increasing the effective intrinsic growth rates of perennials. Plots represent two-species dynamics based on niche overlap (horizontal axis) and species average fitness ratio (vertical axis) between a pair of one annual species and one perennial species. This space is divided into three regions: deterministic exclusion (green), coexistence (blue), and contingent exclusion (orange). The left panel shows the case when perennial life-history processes are not incorporated into the model, while the right panel shows the case when perennial life-history processes are incorporated. Each point represents a pair of species average fitness ratio and niche overlap computed from 2, 000 posterior samples from the posterior distribution of parameter values (the color map represents the density of the points). Note that the species average fitness ratio here refers to the ratio of annual fitness to the perennial fitness, so that the upper green regions correspond to annual-dominated deterministic exclusion and the lower green regions to perennial dominance. Perennial life-history processes only influence the effective intrinsic growth rates, but not the effective competition strength (i.e., life-history processes only change fitness ratios). This implies that including perennial life-history processes increases the proportion of the posterior distribution that falls into the contingent exclusion region (orange region). The details of computing fitness ratio and niche overlap can be found in Appendices A and C, and plots for individual pairs can be found in Appendix E.

**Figure 4:**
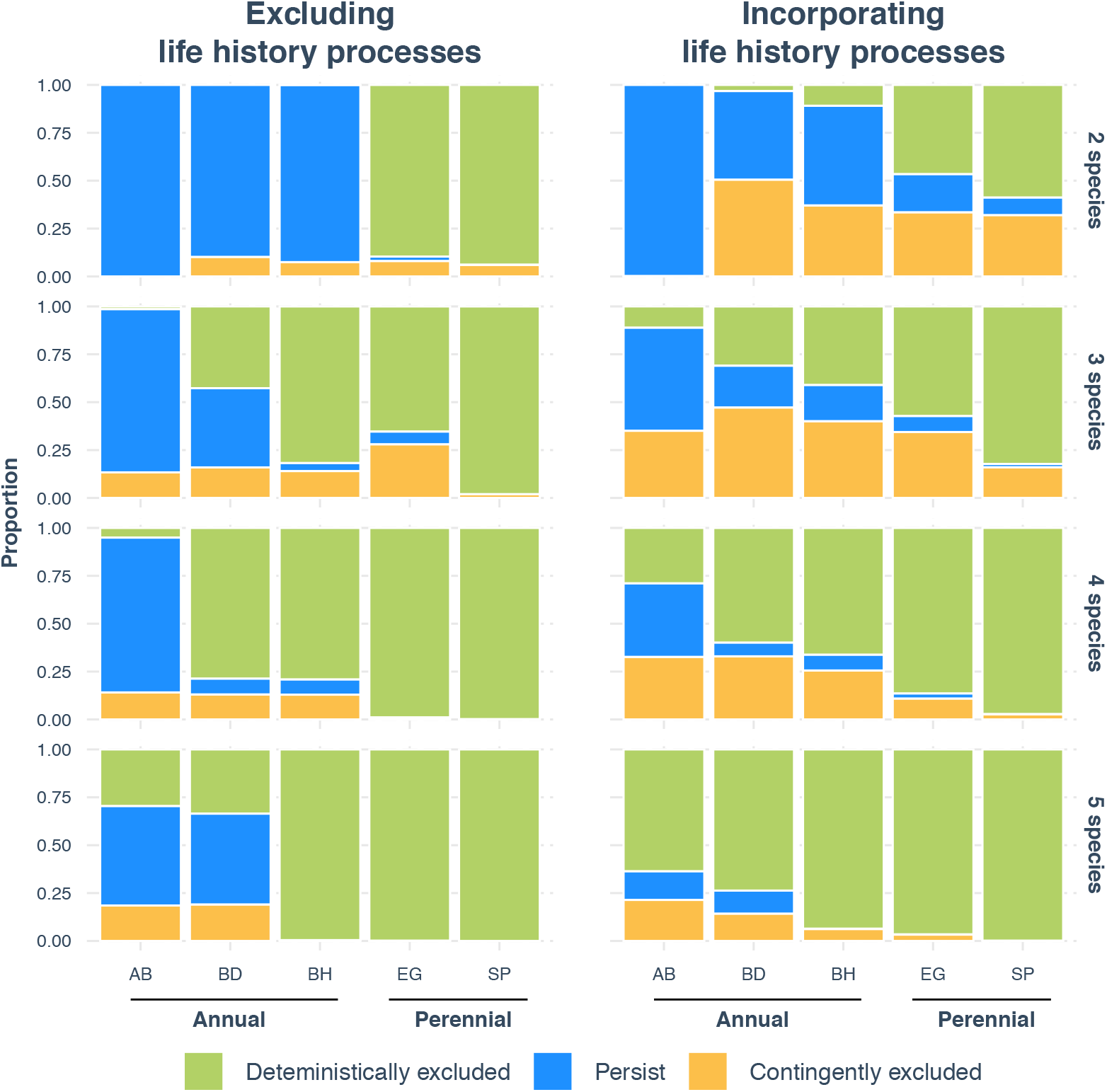
Contingent exclusion is less likely when the community size is larger. We show how the proportions of contingent exclusion, deterministic exclusion, and persistence for each of the five focal species change with community size. The horizontal axis denotes the plant species, where AB stands for *Avena barbata*, BH for *Bromus hordeaceus*, BD for *Bromus diandrus*, EG for *Elymus glaucus*, and SP for *Stipa pulchra*. AB, BD, and BH are annual species while EG and SP are perennial species. We tested all the possible *n*-species combinations with both annual and perennial species present using 2,000 posterior parameter samples. The vertical axis denotes the average proportion of occurrences of deterministic exclusion (green), persistence (blue), or contingent exclusion (orange) in all these combinations. The left and right panels show the case when perennial life-history processes are excluded and included into the model, respectively. The vertical panels show the patterns in each community size (from two-species communities to five-species communities). We found that the proportion of deterministically-excluded species increases with increasing community size (the opposite patterns for contingent exclusion).

Next, we analyzed the effects of community size on the patterns of competitive exclusion. The structural approach argues that contingent exclusion is less likely—and deterministic exclusion more likely—when the community size is larger (Figure 1). Figure 4 confirms this expectation in the empirical data. We found that the percentage of deterministically excluded species rises from 23% in two-species communities to 85% in five-species communities. By contrast, the percentage of contingently excluded species falls from 31% in two-species communities to 9% in five-species communities. Note that we are studying the patterns of competitive exclusion on a species level here (i.e., whether a species persists, is deterministically excluded, or is contingently excluded). In addition, we found that the effect of community size acts more strongly on annual than perennial species (Appendix F). The effect of community size remained consistent with and without incorporating perennial life-history processes (Appendix F).

Lastly, we analyzed the effect of competition structure on the patterns of competitive exclusion. The empirical competition structure (Figure 5A) exhibits two key features: relatively weak intraspecific competition, and intransitive competition. The structural approach establishes that contingent exclusion is more likely when a community has a larger feasibility domain. Figure 5B confirms this expectation in our empirical system: under contingent exclusion, communities have larger feasibility domains (right orange histograms) than the ones generated under deterministic exclusion (left green histograms). Note that the size of feasibility domain decreases as a function of community size, and coexistence (middle blue histograms) is only observed in two-species communities (Fig. 5B). Additionally, we found theoretically (using simulations, as detailed in Methods) that the empirical relationship between competitive exclusion and the size of the feasibility domain emerges by generating weak intraspecific competition structures, regardless of being intransitive or hierarchical (Fig. 5C). These results are robust to different parameterizations in simulations (Appendix G).

**Figure 5:**
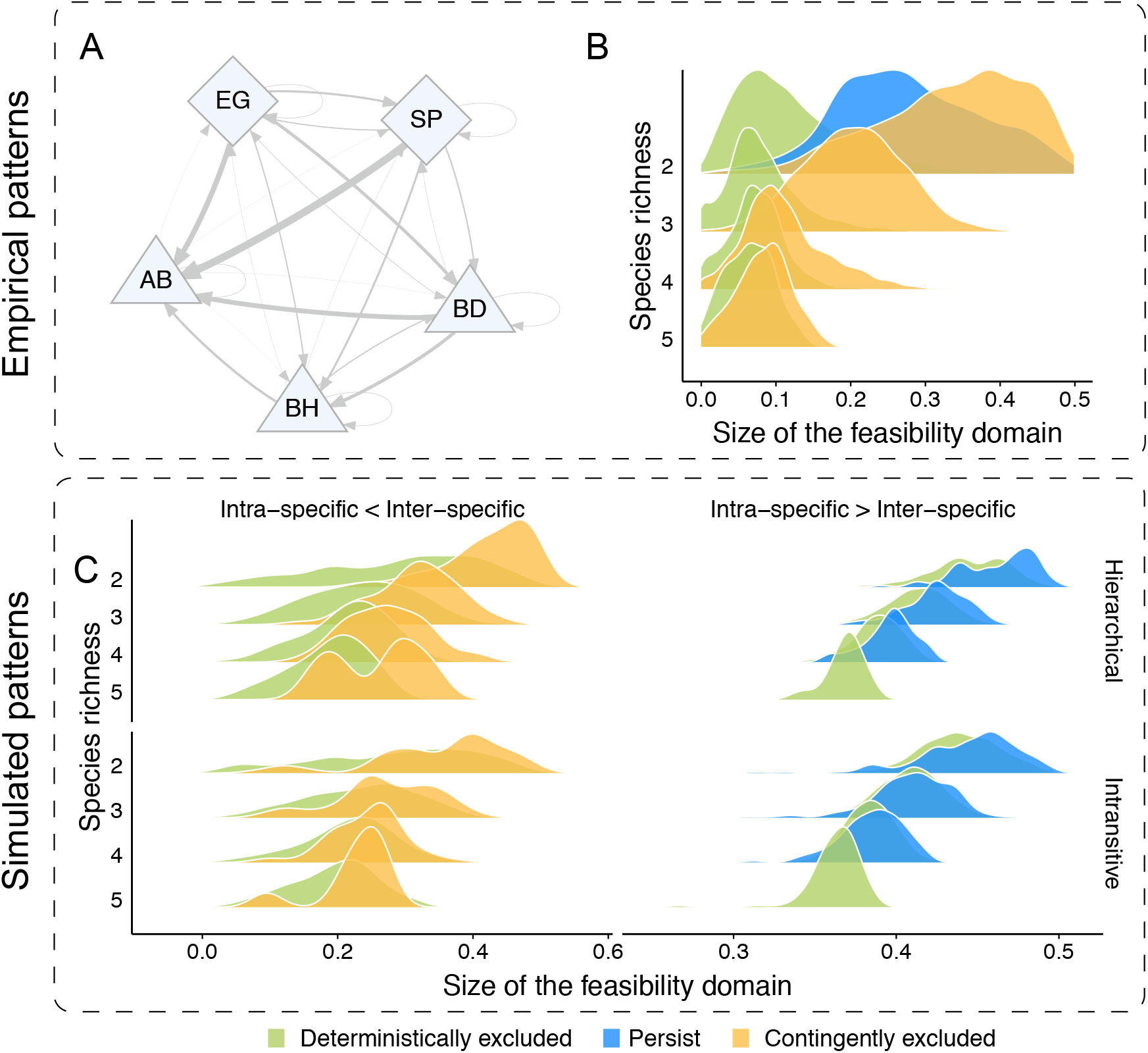
Weak intraspecific and not intransitive competition drives the patterns of competitive exclusion. Panel (**A**) shows the competition structure among annuals and perennials in the empirical data from California grassland plant species. Each node represents a plant species, where the triangles (*Avena barbata* (AB), *Bromus hordeaceus* (BH), and *Bromus diandrus* (BD)) are annuals and the diamonds (*Elymus glaucus* (EG) and *Stipa pulchra* (SP)) are perennials. The direction and width of the links represent the direction and strength (averaged from the posterior samples) of competition. We observe two key structures: (i) intraspecific competition (self-loops) is in general weaker than interspecific competition (edges), and (ii) competition is intransitive (non-hierarchical). Panel (**B**) shows the outcome of competition—deterministically excluded, persist, or contingently excluded—for each empirically-derived parameter set, grouped into histograms by qualitative outcome. We characterize the competition structure of a community across different community sizes using the normalized size of the feasibility domain (horizontal axis). The empirical data show that deterministic exclusion (green histograms) is mostly characterized by structures with a relatively small feasibility domain. Contingent exclusion (orange histograms) have the opposite patterns. Panel (**C**) shows the theoretical expectations about how competition structure affects the patterns of competitive exclusion. We show model-generated communities with different competition structures. We use two structural combinations: (i) communities with either a low (intraspecific < interspecific) or high (intraspecific > interspecific) intraspecific competition, and (ii) communities with either a hierarchical or intransitive competition structure. We find that the competition structures with weaker intraspecific competition, regardless of being hierarchical or not, produce qualitatively the same patterns as the empirical patterns shown in Panel (**B**).

## Discussion

Despite the recent research focus on understanding the mechanisms underlying stable coexistence (Levine & HilleRisLambers, 2009; Adler *et al*., 2007; Chesson, 2000; Godoy *et al*., 2014; Kraft *et al*., 2015), competitive exclusion occurs frequently in nature, and the drivers of deterministic versus contingent exclusion remain poorly understood in multispecies communities (Fukami, 2015; Fukami *et al*., 2016; Uricchio *et al*., 2019; Mordecai *et al*., 2015; Mordecai, 2013). Indeed, in multispecies communities, complex outcomes that combine deterministic and contingent exclusion among groups of species are possible, challenging the extension of results from two-species communities (Case, 1995; Uricchio *et al*., 2019). Here, we provide a theoretical framework following a structural approach to understand the emergence and sources of competitive exclusion in multispecies communities, specifically to distinguish when competitive exclusion is dominated by deterministic or contingent exclusion. We have evaluated three key expectations in multispecies communities derived from our theoretical framework: (i) For contingent exclusion to occur, it is necessary that species have a greater negative effect on their competitor’s per capita growth rate than on their own self-regulation. (ii) The larger the intrinsic growth rates of competitively inferior species, the more likely that contingent exclusion occurs. (iii) The larger the feasibility domain of a community, the more likely that contingent exclusion can be observed. We tested these expectations in an empirical study system composed of five annual and perennial grasses occurring in California grasslands, which exhibit both deterministic and contingent exclusion and several biologically interesting features, including variation in life history strategy, weak self-regulation and strong interspecific competition, and intransitive (non-hierarchical) competition (Uricchio *et al*., 2019). Specifically, we investigated the impact of perennial life-history processes, community size, and competition structure dictate the dynamics of competitive exclusion in this system using the structural approach, which applies to communities larger than two species.

First, we found that perennial life history (interannual survival and reproduction of adult bunchgrasses) increases the probability of observing contingent exclusion by increasing perennial species’ effective intrinsic growth rates (Figures 3 and 4). These life-history processes contribute only to the effective intrinsic growth rates but not to the effective competition strength. In a two-species community, perennial life-history processes increase the fitness of competitively inferior species, making deterministic exclusion less likely (Figure 3). In multispecies communities, we have shown that these life-history processes also help the competitively inferior species (Figure 4). This reveals the importance of life-history processes for increasing the chance of survival of inferior competitors.

Second, we have shown that the probability of observing contingent exclusion decreases with community size (Figure 4). This result is contrary to the naive expectation that contingent exclusion dominates in larger communities, derived from randomly constructed communities (Zhao *et al*., 2020). However, it has remained unclear what happens when communities are structured following a strong deterministic component of population dynamics (Fukami, 2015). For example, in our focal system, annual species are generally superior competitors to perennial species. Under this scenario, contrary to the naive expectation, we should expect to see deterministic exclusion dominating larger communities. That is, a larger community is more likely to contain at least one species that has a large enough competitive advantage over the others to deterministically exclude them. This apparently contradictory expectation aligns well with the intuition derived from our structural approach (Figure 1). Further, these findings reveal that multispecies dynamics may be more predictable than previously thought (May, 1972).

Third, we found that the probability of observing contingent exclusion increases as a function of the size of the feasibility domain defined by the ratio between intraspecific and interspecific competition, and not by the level of hierarchical competition (Figure 5). While many empirical studies have shown that intraspecific competition tends to be stronger than interspecific competition (LaManna *et al*., 2017; Adler *et al*., 2018), recent work has questioned the generality of the empirical evidence supporting stronger intraspecific competition (Hülsmann & Hartig, 2018; Chisholm & Fung, 2018; Detto *et al*., 2019; Broekman *et al*., 2019). Moreover, we have shown that intransitive (or non-hierarchical) competition is unlikely to explain the outcomes of competitive exclusion in the studied system. By contrast, intransitive competition can play an important role in shaping species coexistence (Allesina & Levine, 2011; Soliveres *et al*., 2015; Gallien *et al*., 2017). Thus, our findings imply that ecological mechanisms may play different roles in coexistence and competitive exclusion.

In light of an increasing rate of species invasions as a result of global anthropogenic changes in climate and land use, ecological systems are in dire need of sustainable strategies to mitigate threats to native species. Our study system of grassland plants is an ecologically important and widespread ecosystem that faces such a challenge (Myers *et al*., 2000). It has been suggested that exotic annual grasses have the potential to replace native perennial grasses in over 9 million hectares of California grasslands (Seabloom *et al*., 2003). Indeed, in our study site located in Jasper Ridge Biological Preserve, while these grasses often co-occur at the spatial scale of within *∼*100m of each other, there are many patches where these grasses do not co-occur within *∼*10m. However, given the long time scales for exclusion to fully play out, we cannot say for certain that competitive exclusion would dominate in the system. That is, besides the possibility of competitive exclusion, there are two other possibilities: The first possibility is that a patchwork of different environmental conditions favors different species. For example, we have observed exotic annuals in more disturbed habitats (e.g., *Elymus glaucus* in the zones around oak trees), while native perennials in less distrubed habits (e.g., *Stipa pulchra* in more open grasslands with lower disturbance). The second possibility is that a patchwork of local contingent exclusion dynamics have played out such that species are maintained in local patches that are not truly stably coexisting with other species. Regardless of the specific explanation, this pressing challenge has underscored the need for systematic restoration efforts (Gea-Izquierdo *et al*., 2007; Seabloom, 2011; Werner *et al*., 2016). Our study has shown that the approach to restoration should be different depending on the richness of the system. According to our findings, systems with few species can be strongly driven by contingent exclusion, implying that the restoration can be achieved by focusing on regulating factors, such as life-history traits, self-regulation, or population abundances. By contrast, species-rich systems can be strongly driven by deterministic exclusion, implying that the restoration can be achieved by focusing on limiting factors, such as availability of resources. This result, of course, needs to be taken with caution as we have not used spatialtemporal variation in our analysis (it is empirically challenging to measure local-scale variation in model parameters). This, however, can open a new perspective to restoration management since our key results are testable and generalizable to a wide range of study systems using the same study designs that investigate species coexistence (Levine & HilleRisLambers, 2009; Godoy *et al*., 2014; Adler *et al*., 2018).

Although the understanding of species coexistence has been one of the major topics in ecology for decades (May, 1972; McCann, 2000; Meszéna *et al*., 2006; Ives & Carpenter, 2007; Bastolla *et al*., 2009; Allesina & Tang, 2012; Rohr *et al*., 2014; Barabás *et al*., 2014), competitive exclusion remains the dominant—if hidden—foundation of ecological community structure. While species coexistence and competitive exclusion go hand-in-hand, our understanding about coexistence is much better than exclusion. Competitive exclusion is fundamentally different in two ways: deterministic and contingent. To understand the role of historical contingency in ecological communities, it is paramount to uncover the frequency of and mechanisms underlying deterministic versus contingent exclusion. In this direction, we have taken a new heuristic perspective that partitions exclusion into these two categories within multispecies communities. We hope this work can motivate future research looking into the rich and potentially predictable dynamics of competitive exclusion in multispecies communities.

## Authors’ contributions

All authors conceived the ideas and designed the methodology. C.S. performed the study. S.S. supervised the study. C.S. and S.S. wrote a first version of the manuscript. All authors contributed with substantial revisions. E. M. and L. U. compiled and provided data.

## Data availability

The data of the California grassland community have previously been archived on https://www.journals.uchicago.edu/doi/abs/10.1086/701434. The code supporting our analysis is archived on Github https://github.com/clsong/competitive exclusion.

## Acknowledgement

We thank Mohammad AlAdwani, David L. Des Marais, Lucas P. Medeiros, Caio Guilherme Pereira, and Pengjuan Zu for insightful discussions that led to the improvement of this work.

## Supplementary Material for

## A A brief introduction to Modern Coexistence Theory on competitive exclusion

Modern Coexistence Theory (MCT) is widely adopted to study competitive exclusion (Chesson, 2000; Fukami *et al*., 2016; Ke & Letten, 2018). The canonical formalism of MCT on two-species communities builds upon Lotka-Volterra (LV) competition dynamics. The formulation of two-species LV competition dynamics is written as

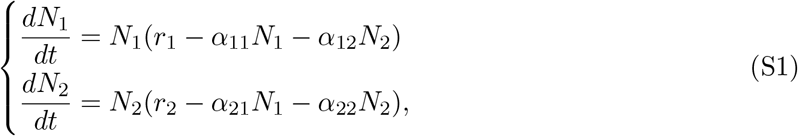

where the variable *N*_*i*_ represents the abundance of species *i*, the parameters *r*_*i*_ > 0 and *α*_*ii*_ > 0 correspond to the intrinsic growth rate and the self-regulation (or intra-specific competition) of species *i*, respectively, and *α*_12_ > 0 and *α*_21_ > 0 are the corresponding interspecific competition strengths.

From the LV competition dynamics, MCT defines niche overlap *ρ* as 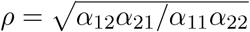, and species average fitness ratio *κ*_2_*/κ*_1_ as 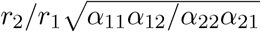 (Chesson, 2018; Bartomeus & Godoy, 2018). Building upon these two concepts, MCT claims that contingent exclusion arises when

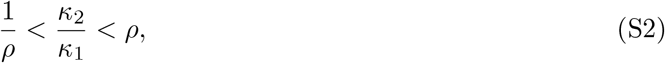

and deterministic exclusion arises when

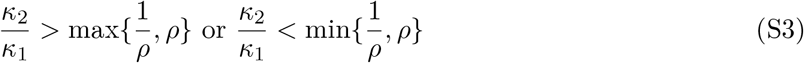

These conditions are illustrated in Figure 3. Note that we used the effective intrinsic growth rates and competition strength in Figure 3 as we translated the population dynamics of grass species into Equation S1.

## B Interpretation of Structural Approach in different theoretical formalisms

The crux of the structural approach is to simplify ecological dynamics as a function of internal and external conditions. In the main text, we have represent external conditions by intrinsic growth rates and represent internal conditions by the competition structure. Here we briefly interpret this representation across several mathematically equivalent but ecologically different theoretical formalism of Lotka-Volterra dynamics. A more detailed discussion can be found in Song *et al*. (2020a).

There are three theoretical formalisms of two-species Lotka-Volterra dynamics. The formalism we adopted in the structural approach (which we call *r*-formalism) is:

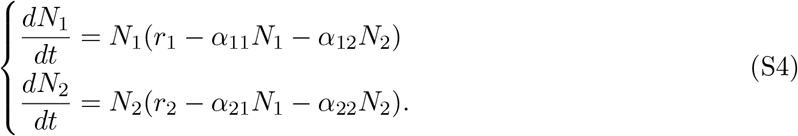

where *r*_*i*_ and *α*_*ij*_ are separated.

Modern Coexistence Theory usually adopts another formalism (which we call MCT-formalism):

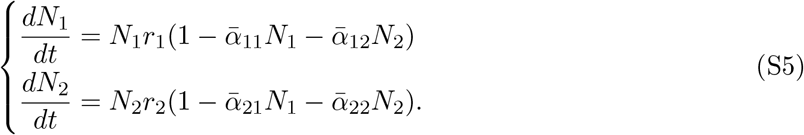

where 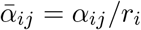. Thus, under the MCT-formalism, *r*_*i*_ and 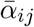 are interlinked. And the third formalism (which we call *K*-formalism) is:

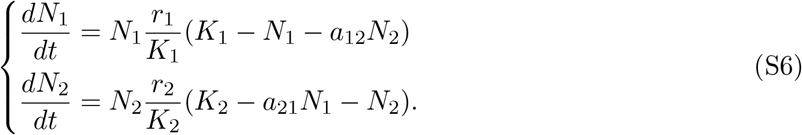

where the competition strength is to be standardized by the intraspecific competition, i.e., *a* _*ij*_ = *α* _*ij*_ */α* _*ii*_.

We first focus on the link between *r*-formalism and MCT-formalism. The ecological interpretations are fundamentally different in these two formalisms. The reason is that while *α*_*ij*_ and 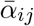 are both called interaction strengths, they have different **units**: *α*_*ij*_ in the *r*-formalism measures the absolute reduction in the growth rates, while 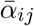 in the MCT-formalism measures the relative reduction in the growth rates to the maximum growth rates. The reason why we have adopted the *r*-formalism is that *α*_*ij*_ in the *r*-formalism is what most empirical studies measure.

We then focus on the link between *r*-formalism and *K*-formalism. To establish the equivalence between the *r*-formalism and the *K*-formalism, the carrying capacity *K*_*i*_ of species *i* and the intrinsic growth rates are linked via *K*_*i*_ = *r*_*i*_*/α*_*ii*_. Thus, if we assume that *α*_*ii*_ is fixed (which is a common assumption in theoretical and empirical studies), then *K*_*i*_ and *r*_*i*_ would reflect identical biotic or abiotic information.

## C Applying the structural approach to the population dynamics of annual and perennial species

### C.1 A brief introduction of the structural approach

Here we present a brief, self-contained description of the structural approach in community ecology. A more detailed, technical description can be found in Song *et al.* (2018b).

Consider an ecological community with *S* interacting species governed by some nonlinear population dynamics. Suppose the equilibrium 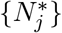 of the community is constrained by a set of linear equations,

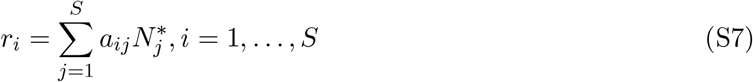

where *r*_*i*_ is referred as the effective intrinsic growth rate and *a*_*ij*_ is referred as the effective interaction strength.

Feasibility of the community refers to the situation in which the equilibrium of all species is positive (i.e., 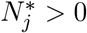, for all *j*) (Roberts, 1974). The feasibility domain *D*_*F*_ —the full set of intrinsic growth rates *r*_*i*_ that gives rise to feasibility—is given by (Logofet, 1993; Song *et al*., 2018b):

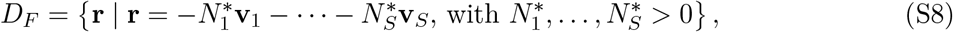

where **v**_*i*_ = {*a*_1*i*_, …, *a*_*Si*_} is the *i*th column vector of the interaction matrix.

Importantly, the operation of positive scalar multiplication on the column space of the effective competition structure **A** does not change the feasibility domain (Song *et al*., 2018b). Specifically, **v**_*i*_ → *c*_*i*_**v**_*i*_ when *c*_*i*_ is some positive constant (equivalently, changing the effective competition strength from *a*_*ij*_ to *c*_*i*_*a*_*ij*_ for all *j*) does not change the feasibility domain.

### C.2 Annual species

We first apply the structural approach to the population dynamics of annual species. As a reminder, the population dynamics of annual species is written as:

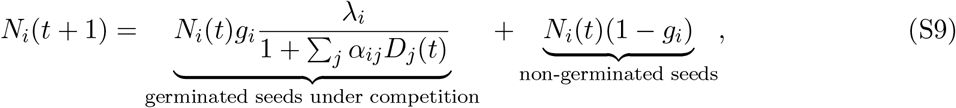

To perform the feasibility analysis in the structural approach, we focus on the equilibrium

*N*_*i*_(*t* + 1) = *N*_*i*_(*t*). The equilibrium condition is equivalent to:

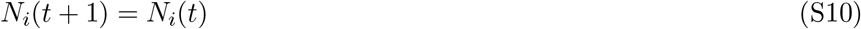

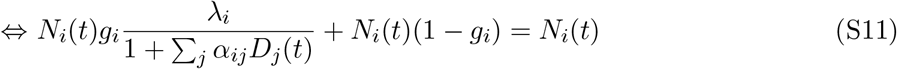

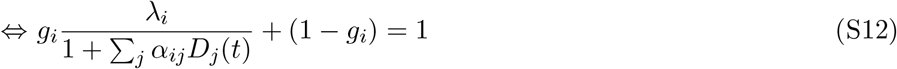

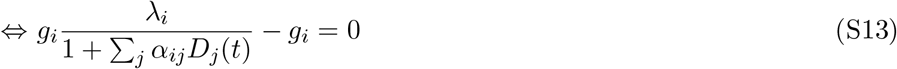

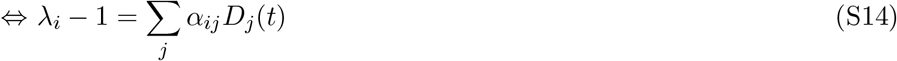

Substituting the definition of *D*_*j*_ from Eqn. (2), the equilibrium condition can be equivalently expressed as:

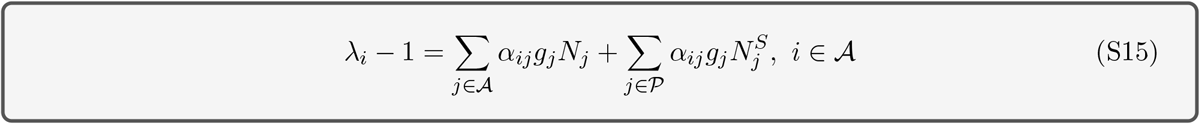

### C.3 Perennial species

Then we apply the structural approach to the population dynamics of perennial species. As a reminder, the population dynamics of perennial species are written as:

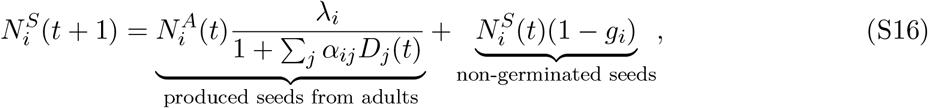

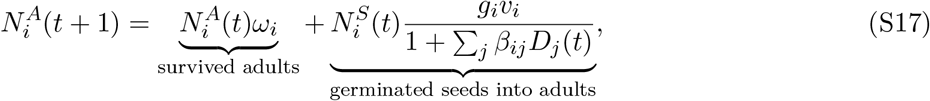

#### C.3.1 Excluding life-history processes in perennial species

When we exclude the life-history processes in perennial species, the equilibrium condition is same as that of annual species (Eqn. S15):

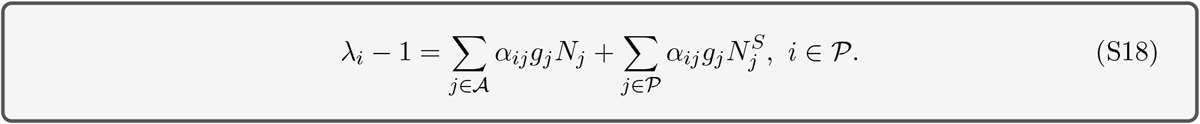

#### C.3.2 Incorporating life-history processes in perennial species

**Without considering the density-dependence in transition from adults to seeds** Here we consider the case when the germinated seeds into adults are not under the pressure of competition. Mathematically, *β*_*ij*_ = 0 in Eqn. 4. Specifically, Eqns. 3 and 4 reduce to:

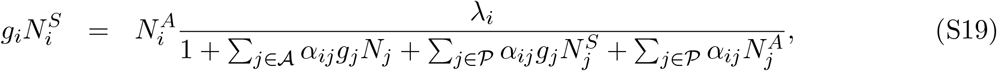

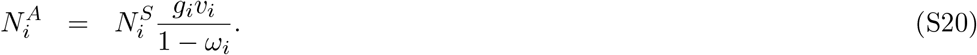

Substituting the expression of 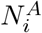 from Eqn. (S20) into Eqn. (S19), the equilibrium conditions are:

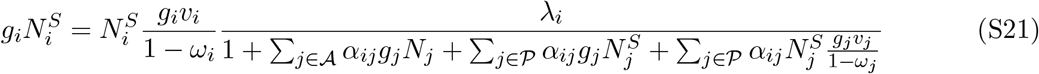

Then the equilibrium condition can be equivalently expressed as:

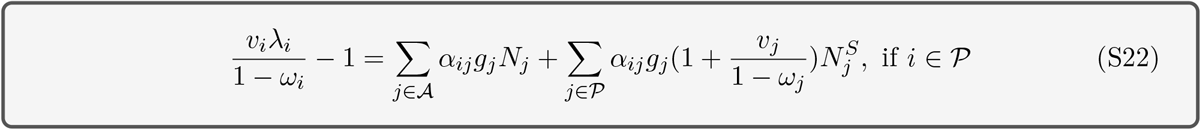

The key difference between Eqn. S18 and S22 is the change of effective parameters:

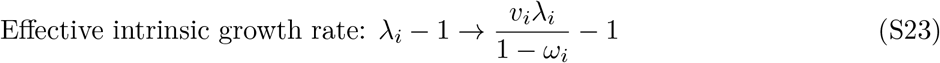

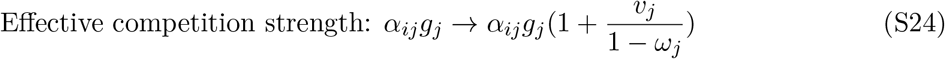

With the effective parameters according to the transformations listed in Eqns. S23 and S24, we would have a system of equations with exactly the same dynamics as the original annual/plant dynamics.

As we have discussed in the beginning of this section, multiplication on the column space of competition strength (*a*_*ij*_ → *c*_*i*_*a*_*ij*_, ∀*j*) does not affect the feasibility domain. Here *c*_*i*_ = 1 for annual species while 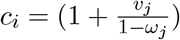 for perennial species. Thus, the feasibility domain remains the same with or without transitions.

Note that this result does not imply that feasibility would not change with or without transitions. As a reminder, the community is feasible if and only if the effective intrinsic growth rates are inside the feasibility domain. Here, the effective intrinsic growth rates changes from *α*_*ij*_*g*_*j*_ to 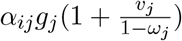 Thus, feasibility (determined by both intrinsic growth rates and competition structure) may change even though the feasibility domain (determined only by the competition structure) does not change.

#### C.3.3 Incorporating life-history processes in perennial species

**Considering the density-dependence in transition from adults to seeds** Here we consider the case when the seeds and adults face the same level of competition. Mathematically, *α*_*ij*_ = *β*_*ij*_. Specifically, Eqns. 3 and 4 reduce to:

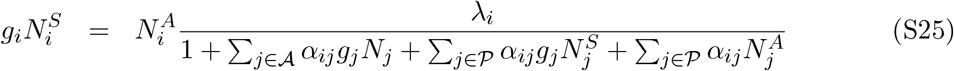

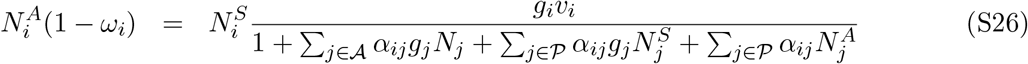

Substituting the expression of 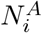 from Eqn. (S26) into Eqn. (S25), the equilibrium conditions are:

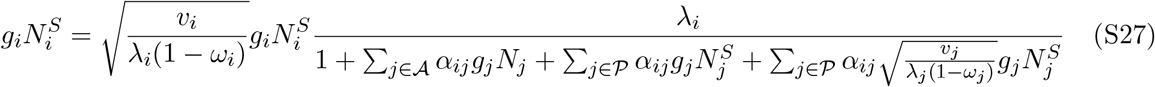

Then the equilibrium condition can be equivalently expressed as:

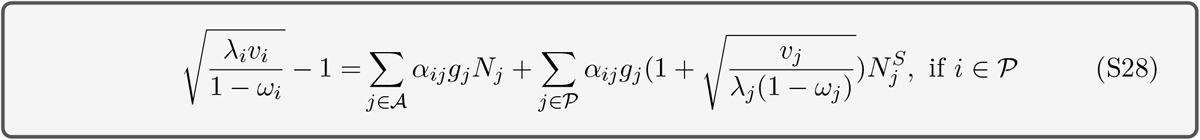

Similarly, we have the changes of effective parameters from Eqn. S18 to Eqn. S28,

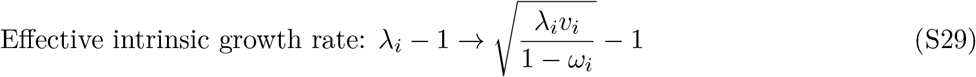

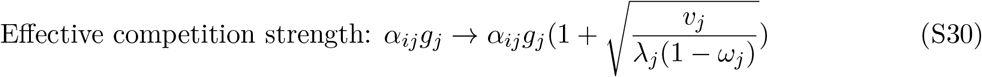

*β*_*ij*_ is the same for all species (i.e., whether *j ∈ 𝒜, 𝒫^S^, 𝒫^A^*)

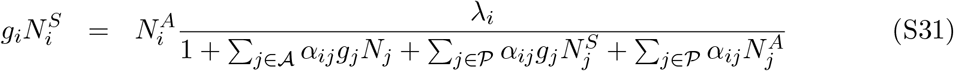

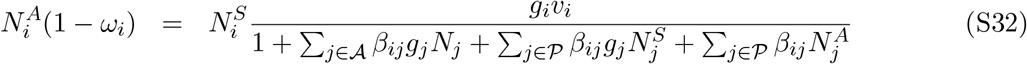

## D Disentangling sources of environmental stress

Here we apply the methods from (Song *et al*., 2020a) to disentangle the effects of parameter perturbations on species pairs. In general, a species pair exhibits a trade-off between the structural stability (tolerance) in competition strength and in intrinsic growth rates. Figure S1 illustrates this trade-off, which is the same for both coexistence and priority effects (Song *et al*., 2020a).

**Figure S1:**
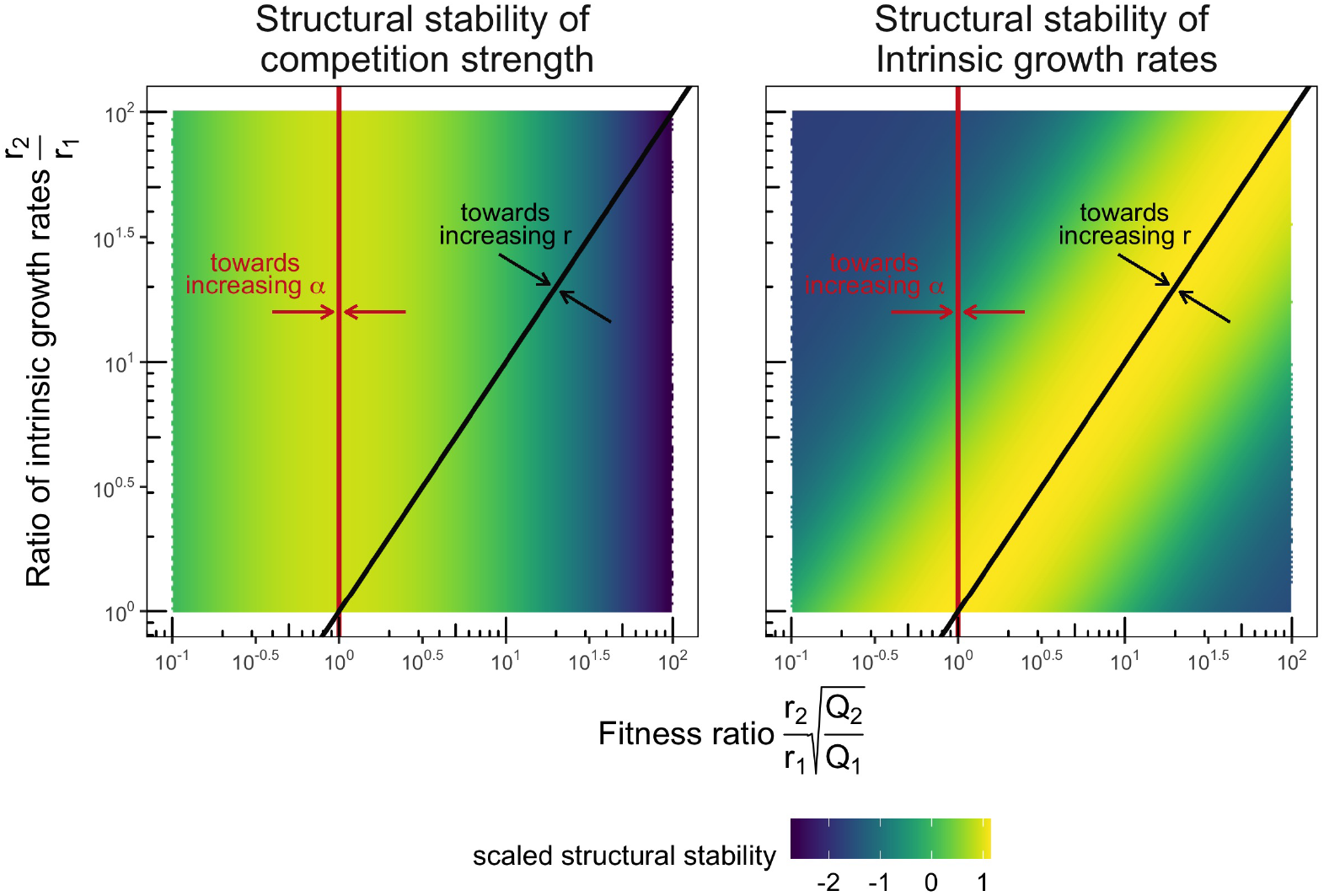
Trade-off between structural stability in competition strength and in intrinsic growth rates. The structural stability in competition strength is increased following the red arrows, and is maximized on the red line (i.e. species average fitness equivalence). The structural stability in intrinsic growth rates is increased following the black arrows, and is maximized on the black line (i.e., species average fitness ratio equals to the ratio of intrinsic growth rates). The color represents the scaled structural stability, where the yellow indicates high while the purple indicates low.

Applying this method to species pairs in the grassland community, Figure S2 shows that: (1) The perennial pairs are robust to both parameter perturbations in intrinsic growth rates and in the competition strength. (2) The annual pairs are more likely to persist under parameter perturbations in the competition strength but not in the intrinsic growth rates. (3) The mixed pairs of one annual and one perennial are robust to changes in intrinsic growth rates only when we exclude the life history processes, but are robust to changes in competition strength only when we incorporate life history processes.

**Figure S2:**
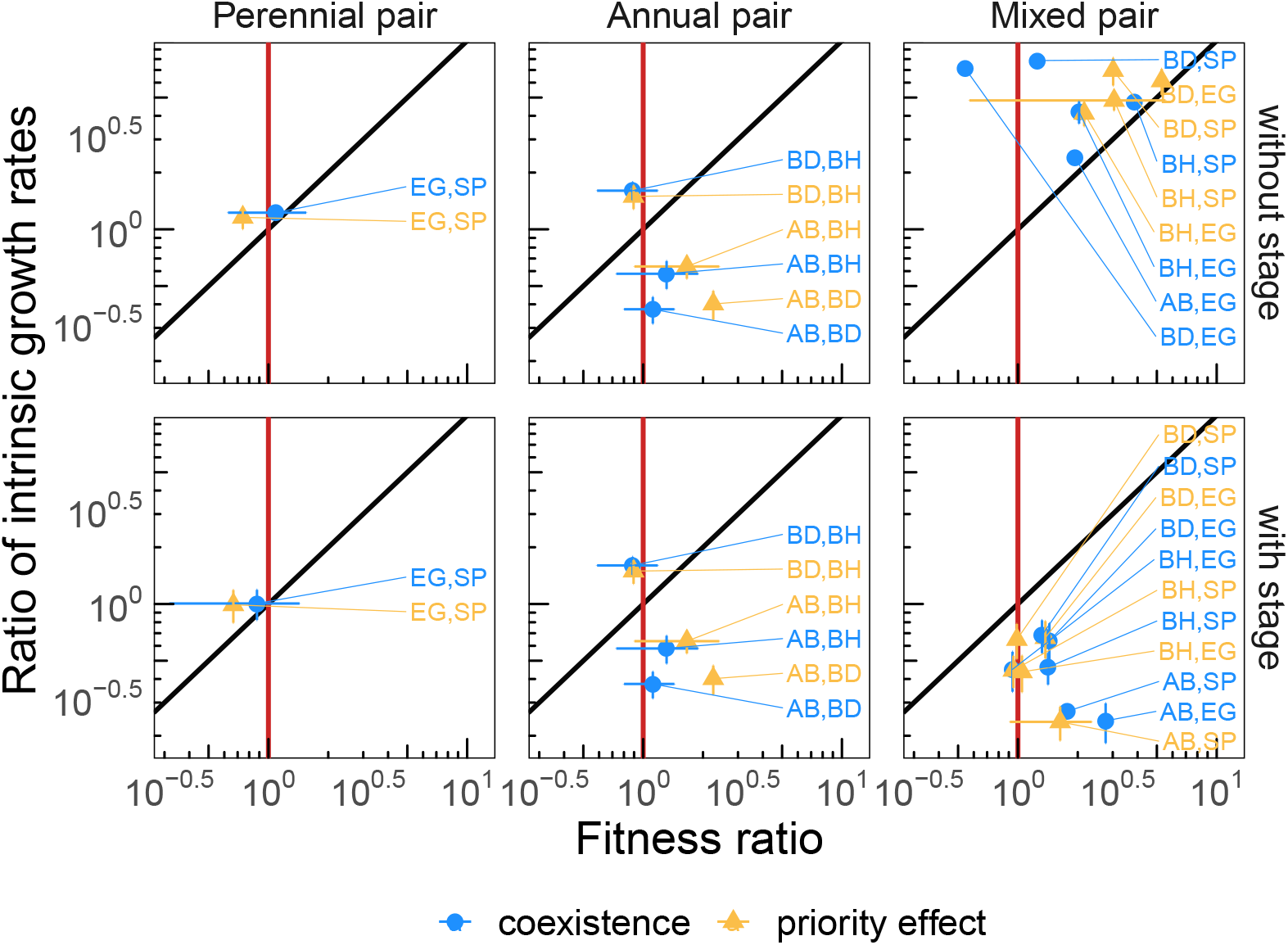
Community persistence under environmental (parameter) perturbations. Here we focus on the structural stability (robustness) of coexistence and priority effects to parameter perturbations. As Figure S1 shows, the structural stability in competition strength increases when the system pair is closer to the red line, while the structural stability in intrinsic growth increases when the system pair is closer to the black line. For the perennial pair (EG & SP; left panels), they maximize both the structural stability in competition strength and in intrinsic growth rates, regardless whether the stage dependency is considered. This result is consistent with the fact that they are native species coexisting for a long time. Then for the annual pairs (middle panels), they tend to maximize the structural stability in competition strength instead of that in intrinsic growth rates. Because the annual species do not have stage dependency, the two panels are exactly the same. Then, for the mixed pairs with one annual and one perennial (right panels), they tend to maximize the structural stability in intrinsic growth rates when the stage dependency is not considered (top), while they maximize the structural stability in competition strength when the stage dependency is considered (bottom). Thus, the stage dependency makes the perennials more vulnerable to parameter perturbations in competition strength (while the annuals have been adapted to these kinds of perturbations). The blue dots denote the pairs exhibiting priority effects, while the orange triangles denote the pairs exhibiting priority effects. The error bars represent two standard deviations.

## E Effects of life-history processes

Figure S3 is a remake of Figure 3 in the main text except that all the species pairs are shown individually..

**Figure S3:**
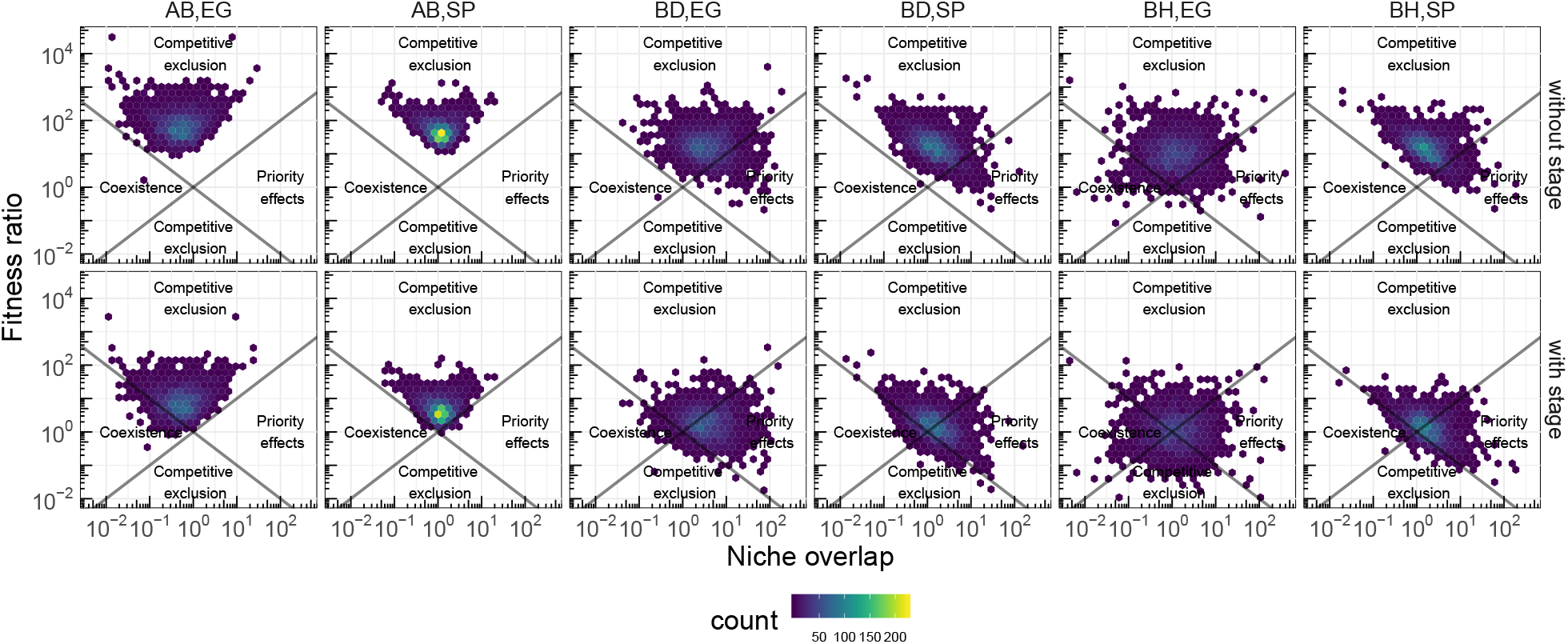
This figure is identical to Figure 3 except species pairs are shown separately

Figure S4 shows the transition probability of community dynamics for a given ecological community between excluding and incorporating perennial life history processes. Note that there is zero transition probability from coexistence to contingent exclusion. The reason is that changing the effective intrinsic growth rates cannot change the system from coexistence to priority effect, or vice versa (Song *et al*., 2020a).

**Figure S4:**
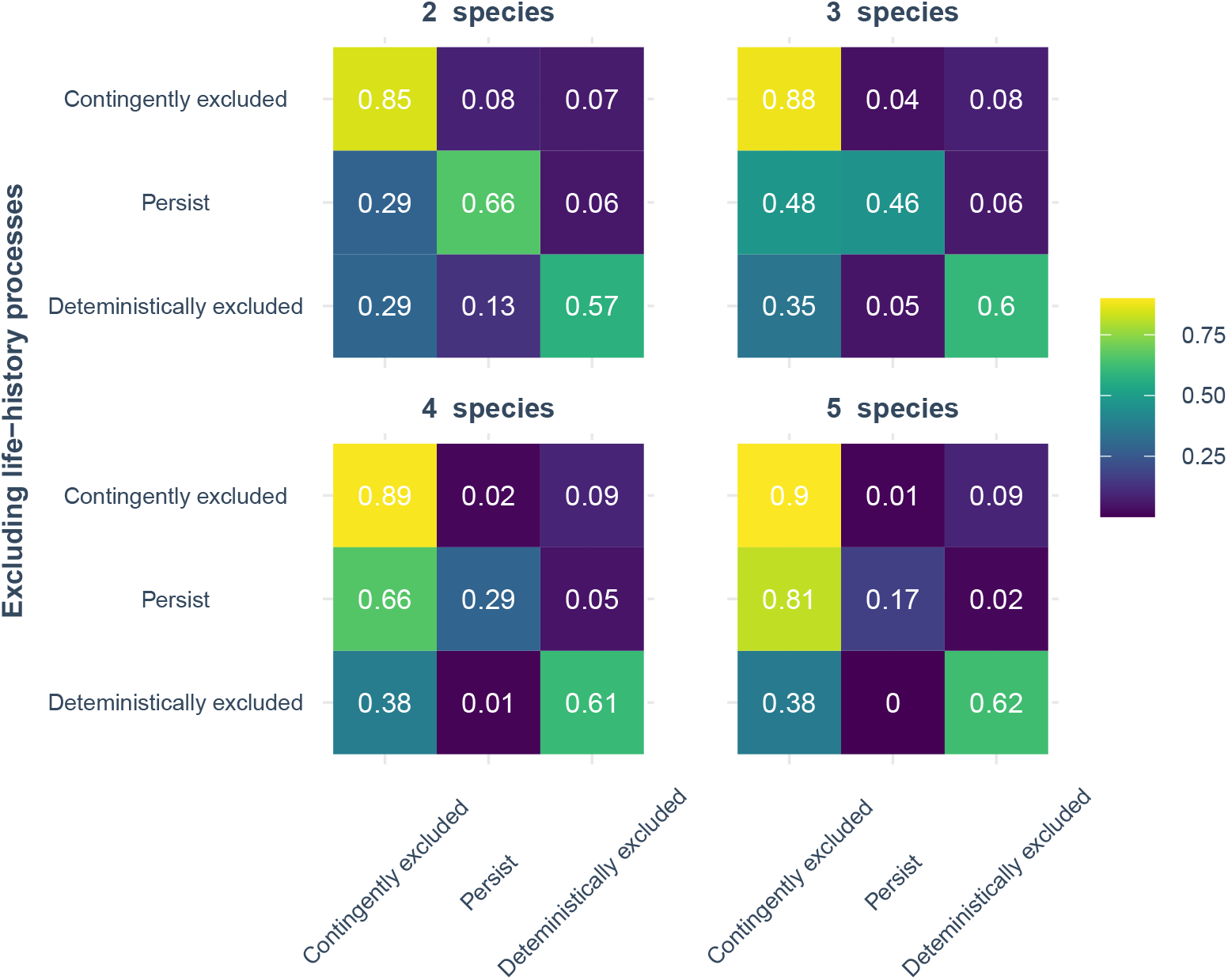
The frequency and prevalence of contingent exclusion decreases as a function of community size. We show the transition matrix of community dynamics between excluding (rows) and including (columns) life-history processes as a function of community size. Each element corresponds to the conditional probability (expressed as frequency) of having a particular dynamics by incorporating life-history processes (e.g., contingent exclusion including life-history, first column) given that the system started in a given dynamics excluding life-history processes (deterministic exclusion, third row). The matrices show that the prevalence (starting and remaining) of contingent exclusion (first element) decreases in general with community size. The matrices also show that the incidence (starting from deterministic exclusion—note that coexistence never leads to contingent exclusion) of contingent exclusion also decreases with community size.

## F Effects of community size

**Figure S5:**
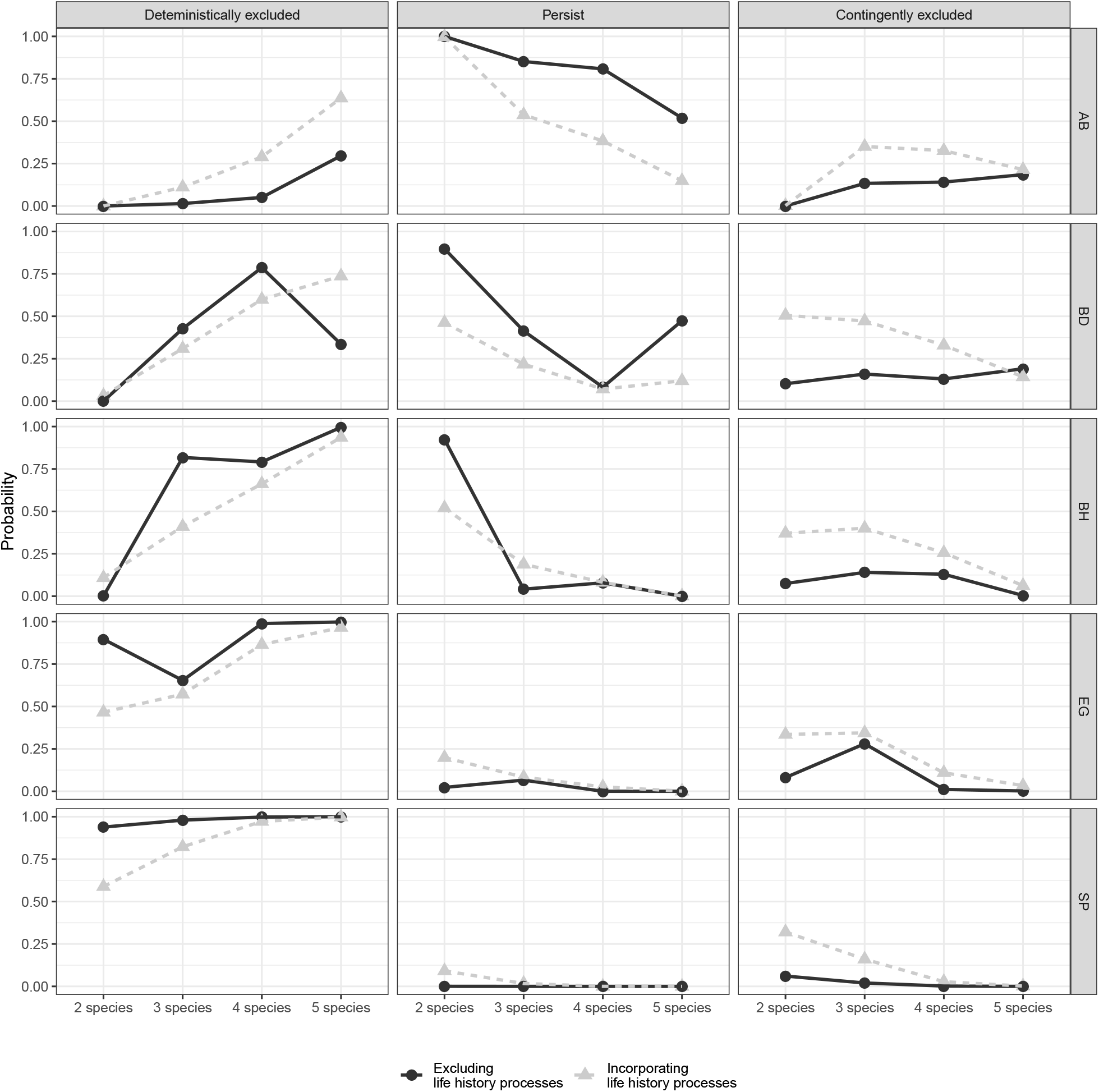
This figure is a remake of Figure 4 in the main text except that the probability is now shown in a scatter plot instead of in a bar plot.

## G Effects of competition structure

Here we perform additional simulations to test the robustness of Figure 5.

We changed the distribution of inter-specific interaction from uniform distribution to half-normal distribution (1*N* (0, 1)1). See Figure S6.

**Figure S6:**
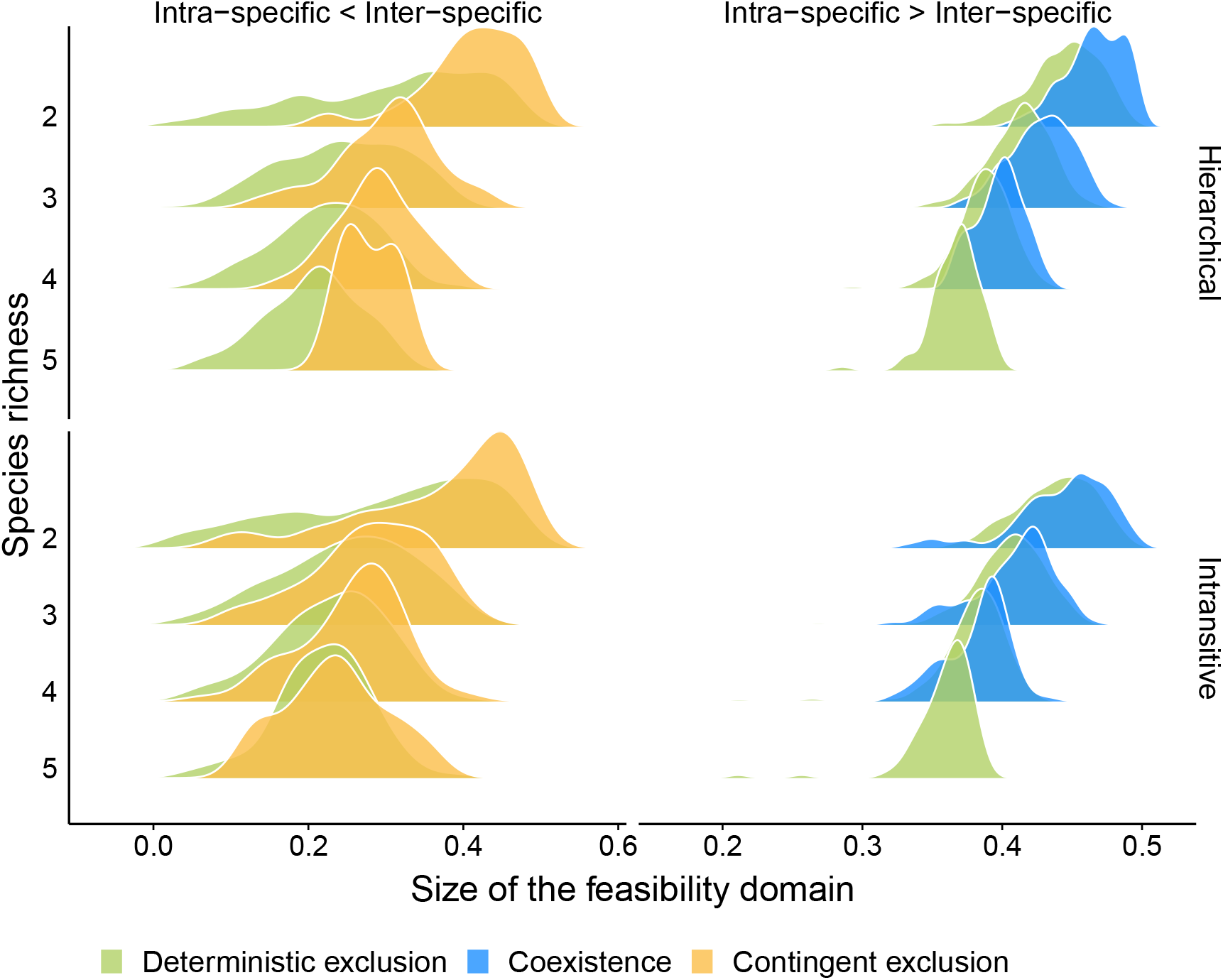
Same as Figure 5 except that the interspecific interactions are drawn from a half-normal distribution. Specifically, this figure shows the theoretical expectations about how competition structure affects the patterns of competitive exclusion. We show model-generated communities with different competition structures. We use two structural combinations: (i) communities with either a low (intraspecific < interspecific) or high (intraspecific > interspecific) intraspecific competition, and (ii) communities with either a hierarchical or intransitive competition structure. We find that the competition structures with weaker intraspecific competition, regardless of being hierarchical or not, produce qualitatively the same patterns as the empirical patterns shown in Panel (**B**) in Figure 5.

## Notes

### Competing Interest Statement

The authors have declared no competing interest.

### Summary of Updates

We have improved the exposition of our methodology and expanding the discussions of the implications to conservation management.

